# Detergent lysis controls are critical for single extracellular vesicle phenotyping in SP-IRIS platforms

**DOI:** 10.64898/2026.10.23.760472

**Authors:** Gonzalo Germán Guendulain, David Otaegui, Hirune Crespillo-Velasco, David Campo-Caballero, Rocío del Carmen Bravo-Miana, Ana Lis Moyano

## Abstract

Bulk analytical methods are extensively used to identify molecular cargo in extracellular vesicles (EVs), but they cannot distinguish nanovesicles from co-isolated non-vesicular particles or resolve discrete EV subpopulations. Single-particle interferometric reflectance imaging coupled with immunofluorescence (SP-IRIS/IF) has emerged as a platform to simultaneously resolve particle size, count, and cargo at single-EV resolution. However, differentiating authentic EVs from non-vesicular signals requires rigorous methodological controls that remain widely underreported in standard workflows. In this study, we analyzed human neural secretomes and serum EVs across different SP-IRIS/IF setups to evaluate their analytical specificity. By using detergent-mediated lysis controls, we found a critical technical limitation in standard single-particle analysis: the persistence of both interferometric and fluorescent signals after detergent treatment in a subset of presumed single-EV events that were non-vesicular artifacts. These findings indicate that lacking appropriate controls on single-EV platforms risks data misinterpretation, underscoring the indispensable implementation of detergent-mediated lysis for single-particle analysis in SP-IRIS/IF platforms.

## Introduction

Extracellular vesicles (EVs) are nanoscale, lipid bilayer-delimited particles that mediate intercellular communication by transporting bioactive cargo, including proteins, lipids, and diverse RNA biotypes. Because EVs can cross biological barriers and circulate systemically through biofluids such as blood, they might mirror the physiological and pathological state of their tissue of origin (Bravo Miana et al., 2024; Kalluri & LeBleu, 2020). Beyond their cargo composition, the physicochemical properties and structural integrity of EVs convey critical information, driving significant interest in their clinical translation. Consequently, high-resolution technical approaches to accurately quantify EV size, morphology, and the physicochemical and molecular properties at single-particle resolution are essential for advancing both basic EV biology and translational biomarker discovery (Bordanaba-Florit et al., 2021; Wang et al., 2022).

Conventional characterization approaches rely primarily on bulk measurements (*e.g.*, Western blot, ELISA, and mass spectrometry), which intrinsically average distinct molecular signatures across heterogeneous vesicle populations. Consequently, discrete EV subpopulations are masked, obscuring molecular patterns that can only be resolved at the single-EV level (Su et al., 2025). To address this limitation, an expanding repertoire of optical, microfluidics and sensor-based technologies have emerged, such as nano-flow cytometry, super-resolution microscopy and single-particle interferometric reflectance imaging sensing (SP-IRIS) (Alexandre et al., 2025; Bravo Miana et al., 2024; Fagúndez et al., 2025; Nevo et al., 2026). However, studying particles at the nanoscale (specifically small EVs < 200 nm) introduces substantial analytical challenges, demanding rigorous assay controls to differentiate genuine EVs from co-isolated non-vesicular macromolecular complexes, lipoproteins, and technical noise (Welsh et al., 2024).

SP-IRIS coupled with immunofluorescence (SP-IRIS/IF) relies on the selective capture of EVs by immobilized antibodies, where single-vesicle size and count are quantified via optical interferometry, while the molecular phenotype of captured EVs is profiled using fluorescently labeled detection antibodies. Although the Minimal Information for Studies of Extracellular Vesicles (MISEV) guidelines suggest essential biochemical controls for single-EV platforms (Welsh et al., 2024), most remain widely underreported in both commercial and custom SP-IRIS/IF workflows. Here, we show that detergent lysis is a critical control to validate SP-IRIS/IF data in single-EV detection and molecular characterization. As proof of concept, we analyzed human neural secretomes and serum across different commercial chips, identifying non-vesicular artifacts with interferometric and fluorescent signals that mimic true EV parameters. Collectively, our findings demonstrate that detergent-lysed controls are critical for recognizing authentic EV detection from non-vesicular particles or antibody-mediated artifacts in SP-IRIS/IF, highlighting the importance of standardized quality-control strategies for single-EV profiling platforms.

## Materials and methods

### Stem cell culture and media collection

Two human induced pluripotent stem cell (hiPSC) clones (F2A112 and F2A121) were previously reprogrammed from a healthy donor with written informed consent and characterized under institutional ethical approval (Herrera Lopez et al., 2025; Wilson et al., 2026). hiPSCs were cultured on Matrigel-coated cultureware (Corning Life Sciences, #354234) in mTeSR™ Plus medium (STEMCELL Technologies, #100-0276), fed every other day, and passaged as small aggregates every 6 to 7 days. Neural induction and human neural rosette (hNR) generation were performed using an embryoid body (EB)-based differentiation protocol. Briefly, hiPSCs were dissociated into a single-cell suspension using Gentle Cell Dissociation Reagent (STEMCELL, #100-0485) and resuspended in STEMdiff™ Neural Induction Medium with SMADi (STEMCELL, #08581), supplemented with RevitaCell (Thermo Fisher Scientific, #A26445-01). To generate uniform EBs, cells were seeded into AggreWell™ 800 plates (STEMCELL, #34811) at a density calculated to yield 3,000 cells per EB. Plates were centrifuged at 100 ×*g* for 3 min to capture cells and incubated at 37 °C. After 24 h, RevitaCell was withdrawn, and partial media changes (3/4 of total volume) were performed daily for 4 days. EBs were harvested, and 300 EBs were plated per well onto Matrigel-coated 6-well plates in STEMdiff™ Neural Induction Medium. Cultures were maintained with daily total media changes. Between days 8 and 11 of differentiation, upon the establishment of hNRs morphology, serum-free conditioned medium (CM) was collected daily (2 mL per well). The collected CM were centrifuged for 30 min at 2,000 ×*g* at 4 °C, filtered through a 0.22 μm PES membrane, aliquoted, and immediately stored at −80 °C until further use.

### Immunocytochemistry and confocal microscopy

Cells were plated onto chambered coverslip (µ-Slide 8-well, IBIDI, #80821) precoated with Matrigel and fixed with 4% paraformaldehyde (PFA) at room temperature (RT) for 15 min. After 3 washes with phosphate-buffered saline (PBS), cells were permeabilized with 0.2% Triton X-100 in PBS. Subsequently, samples were incubated overnight at 4 °C with specific combinations of the following primary antibodies in PBS containing 0.1% Tween-20 (PBST): rabbit anti-SOX2 (1:400, Abcam, #ab97959), mouse anti-α-Tubulin (1:1500, Cell Signaling, #3873), mouse anti-Nestin (1:200, Abcam, #ab22035), and rat AA3 monoclonal hybridoma against the mayor proteolipid protein (PLP1, 1:20). The AA3 antibody was a kind gift from Dr. Irene Givogri and Dr. Ernesto Bongarzone. After 3 washes with PBST at RT, cells were incubated at RT for 1 h with the appropriate secondary antibodies: anti-mouse IgG (1:1000, Alexa Fluor 488 or 647, Invitrogen, #A21202 or #A31571), anti-rabbit IgG (1:1000, Alexa Fluor 488 or Alexa Fluor Plus 555; Invitrogen, #A21206 or #A32794) and anti-rat IgG (1:800, Invitrogen, #A21247). After 3 washes with PBST, cells were mounted using mounting medium containing DAPI (IBIDI, #50011). Images were acquired using a Zeiss LSM 900 Confocal Optical Microscope at Biogipuzkoa Health Research Institute and analyzed using ImageJ software (NIH, Bethesda, MD).

### Analysis of hNR secretomes by SP-IRIS/IF

CM was diluted 1:10 in incubation buffer (IB, provided by the chip manufacturer), and 50 μL of the dilution was incubated on human tetraspanin chips (Leprechaun Exosome Human Tetraspanin Kit, Unchained Labs, #251-1044) for 16 h at RT. These chips are functionalized with capture antibodies targeting human CD9 (clone HI9a), CD63 (clone H5C6), CD81 (clone JS-81), and a mouse IgG isotype control (clone MOPC-21). After incubation, chips were processed using the automated “Exosome Cargo” protocol in the manufacturer’s chip washer (Unchained Labs). To allow intra-vesicular staining, captured EVs were permeabilized for 20 min using a Tween-based buffer (Solution A, provided in the kit) and incubated for 1 h at RT with fluorescently labelled detection antibodies against myelin proteolipid protein (PLP1)-AF® 647 (Novus Biologicals, #NBP3-08534AF647), syntenin-1 (Syn)-CF® 555 (Leprechaun Exosome Cargo Upgrade, #251-1049), and CD9-CF® 488A (provided in the Exosome Human Tetraspanin kit). Chips were dried for 30 min at RT and imaged using the SP-IRIS ExoView R200+ platform with ExoView scanner software v3.2.1 (Unchained Labs). For detergent lysis controls, Triton X-100 was added to the diluted CM to a final concentration of 1%, and incubated for 30 min at RT. Following centrifugation at 10,000 ×g for 15 min, supernatants were incubated on human tetraspanin chips for 16 h at RT. All subsequent processing and imaging steps were performed as described above.

### Human sera collection and processing

Blood samples were collected from healthy controls via venipuncture for diagnostic purposes. This study was approved by the Basque Ethics Committee for Research with Medicines (DOB-EVA-2025-01) on 18 November 2025, in accordance with the Declaration of Helsinki and with written informed consent signed by all participants. Whole blood was collected in BD Vacutainer SST II Advance (Becton, Dickinson and Company, #367953) and allowed to clot at RT for **30** min. Sera was separated by centrifugation at 1500 ×*g* for **10** min at **RT**. The remaining aliquots were processed and immediately stored at −80 °C at the Basque Biobank (BIOEF, Donostia University Hospital, San Sebastián, Spain). Samples were thawed on ice and centrifuged at 2,500 ×*g* for 15 min at 4 °C before use.

### Analysis of human sera by SP-IRIS/IF

Human serum analysis was performed as previously described (Bravo-Miana et al., 2024). Briefly, samples were diluted 1:10 in IB and 50 μL of the dilution were incubated for 2 h at RT on ExoView Human Plasma chips (Leprechaun Exosome Human Plasma Kit, Unchained Labs, #251-1045). In addition to the previously mentioned tetraspanins (Exosome Human Tetraspanin Kit), these chips include capture spots for CD41a+ platelet-derived EVs (clone HIP8). ExoView FLEX chips (Leprechaun Exosome Human Tetraspanin Flex Kit, Unchained Labs, #251-1047) were customized using a monoclonal antibody recognizing an extracellular loop of PLP1 (clone 2D7, Abnova, #H00005354-M03), following manufacturer’s recommendation. Due to the inherent complexity of serum samples, a customized version of the Exosome Cargo protocol was implemented. Briefly, the standard washing sequence was modified to include 6 cycles with Solution A (1,000 μL/well), followed by 8 cycles with Solution B (1,500 μL/well) and 15 cycles with distilled water (2,000 μL/well). EVs staining was performed using fluorescently labelled detection antibodies against PLP1-AF® 647, Syn-CF® 555 or CD81-CF® 555, and CD9-CF® 488A (provided in the kit). Chips were dried for 30 min at RT and imaged as described for the CM analysis. Detergent lysis controls were performed as in CM. After centrifugation, samples were incubated on human plasma chips for 2 h at RT. All subsequent processing and imaging steps were performed as described above for serum samples.

### Data acquisition, processing, and background correction

Data were analyzed using NanoViewer 2.8.10 Software (NanoView Biosciences). Fluorescence intensity cut-offs were defined to filter out non-specific background. For each detection antibody, cut-off values were set so that between 10-20% of the events on the MIgG1 spots were counted as positive. For the CM analysis on human tetraspanin chips, cut-offs ranges ([minimun - maximum]) were set at [350-400 to 65536], [250-300 to 65536], and [350 to 65536] for PLP1, Syn, and CD9, respectively. For serum analysis on ExoView Human Plasma chips and ExoView Flex chips, cut-offs were set at [350-500 to 65536], [250-300 to 65536] and [350-400 to 65536] for PLP1, CD81 and CD9, respectively. After applying these thresholds, any remaining positive events on the MIgG1 spots were considered as basal background. Finally, data from all specific capture spots was normalized by subtracting this background. Mean fluorescence intensity analyses were performed by only considering events with a fluorescence intensity greater than the mean plus three standard deviations (mean + 3 SD) of the MIgG1 for each detection antibody.

## Results

### Cellular model characterization and SP-IRIS/IF setup

Human-induced pluripotent stem cells (hiPSCs) were differentiated into neural rosettes (hNRs) and characterized by immunofluorescence (Figure 1A-B). Undifferentiated hiPSC colonies displayed characteristic compact morphology and expressed the pluripotency transcription factor SOX2. Upon differentiation, hNRs were radially organized and expressed the neural stem cell markers Nestin and SOX2 and the Proteolipid Protein 1 (PLP1) (Figure 1B). PLP1 is an abundant transmembrane myelin protein recently identified by mass spectrometry as a molecular cargo of hNR derived EVs (hNR-EVs), with a potential role in neurite outgrowth and neuron maturation in both murine and human *in vitro* models (Herrera Lopez et al., 2025). Based on this background, our goal was to systematically characterize PLP1 distribution at single-EV resolution. Therefore, we implemented SP-IRIS/IF (ExoView platform), which enables simultaneous sizing, count and multiplexed cargo phenotyping. SP-IRIS/IF setup included sensor chips displaying capture spots functionalized with antibodies targeting EV tetraspanins (CD63, CD9 and CD81), and a mouse isotype control (MIgG1) to assess non-specific binding. Captured vesicles were subsequently probed with fluorescently labelled detection antibodies against CD9, syntenin-1 (Syn) and PLP1 to evaluate their distribution in single EVs (Figure 1C-D).

**Figure 1.**
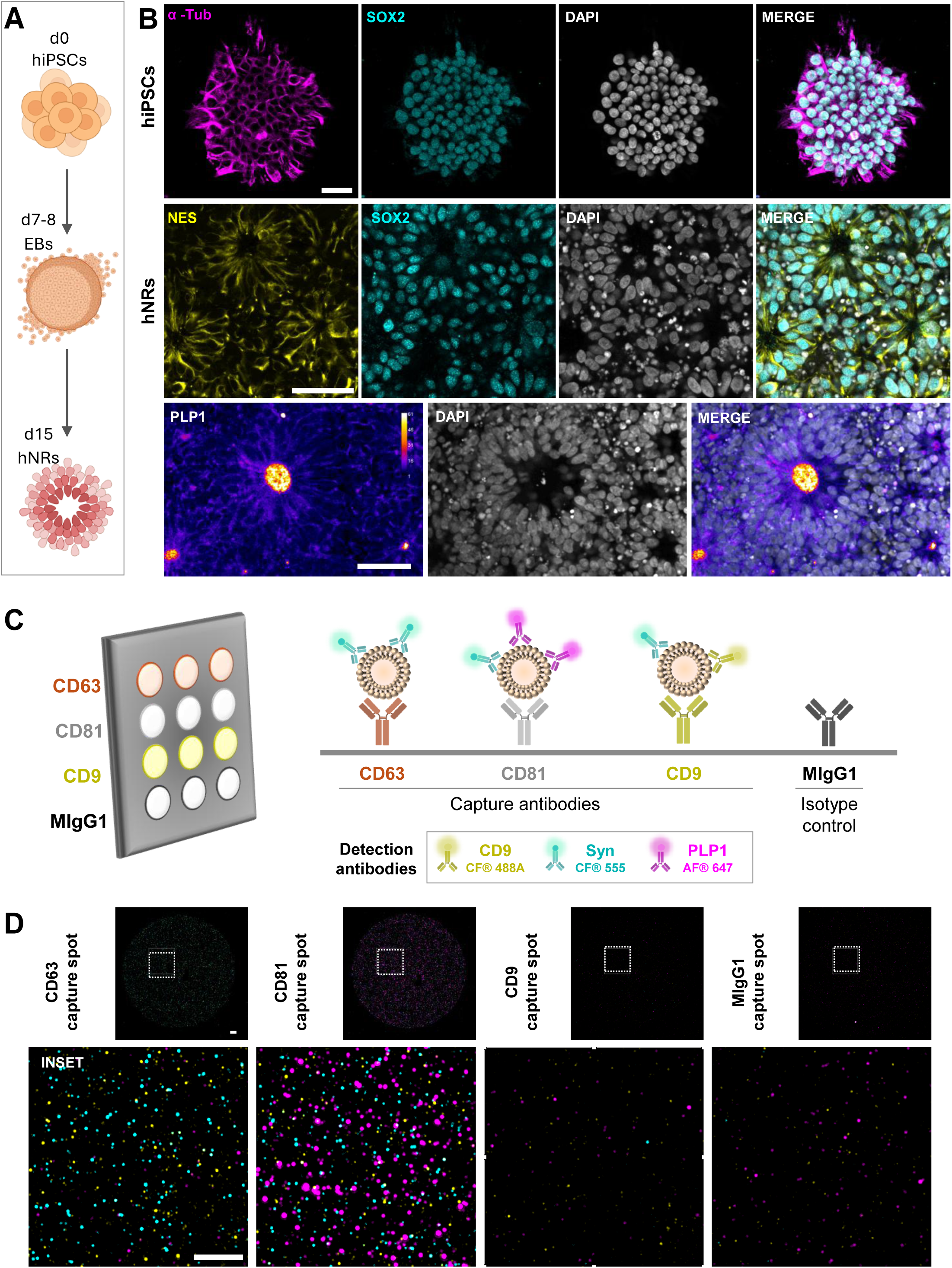
Human neural rosettes culture and SP-IRIS/IF setup. (**A**) Timeline depicting human-induced pluripotent stem cells (hiPSCs; day 0) aggregating into embryoid bodies (EBs; days 7–8) and maturing into radially organized human neural rosettes (hNRs; day 15). (**B**) Representative confocal micrographs of undifferentiated hiPSC colonies (top row) immunostained for α-tubulin (α-Tub, magenta), SOX2 (cyan). Middle row: hNR cytoarchitecture co-expressing the neural stem cell marker Nestin (NES, yellow) and SOX2 (cyan). Bottom row: radial distribution of Proteolipid Protein 1 (PLP1, Fire pseudo-color scale). Nuclei stained with DAPI (gray). Scale bar: 50 μm. (**C**) Schematic SP-IRIS/IF sensor array: sensor chip with capture antibodies targeting tetraspanins (CD63, CD81, and CD9) and a mouse isotype control (MIgG). Captured EVs are analyzed using fluorescent detection antibodies: anti-CD9 CF® 488 (yellow), anti-Syntenin-1 (Syn) CF® 555 (cyan), and anti-PLP1 AF® 647 (magenta). (**D**) Representative micrographs showing single-EV events using detection antibodies against CD9 (yellow), Syn (cyan) and PLP1 (magenta) in capture spots of human neural rosette conditioned medium (hNR-CM). Dashed lines in capture spots delineate insets. Scale bar: 10 μm.

PLP1 is a four-pass transmembrane protein structurally similar to tetraspanins. To determine its association to the captured EVs, we used a detection antibody recognizing an intraluminal epitope of PLP1 (not previously validated for SP-IRIS/IF). To evaluate fixation and permeabilization conditions we implemented manufacturer-validated detection antibodies against Syn (soluble EV cargo) and CD9 (transmembrane EV cargo) (Figure 2A). We compared PBS, Solution A (Tween-based solution provided by the manufacturer), and a mixture of Solutions C and D (fixation and permeabilization solutions, respectively) included in the manufacturer’s kit recommended for intraluminal EV cargo detection. Solution A yielded the highest relative counts of PLP1+ fluorescent events, with only slightly lower Syn+ counts than Solutions C and D, while maintaining CD9+ counts near to PBS control levels (Figure 2B). These results helped us to establish Solution A as the optimal condition to characterize PLP1 distribution in captured EVs.

**Figure 2.**
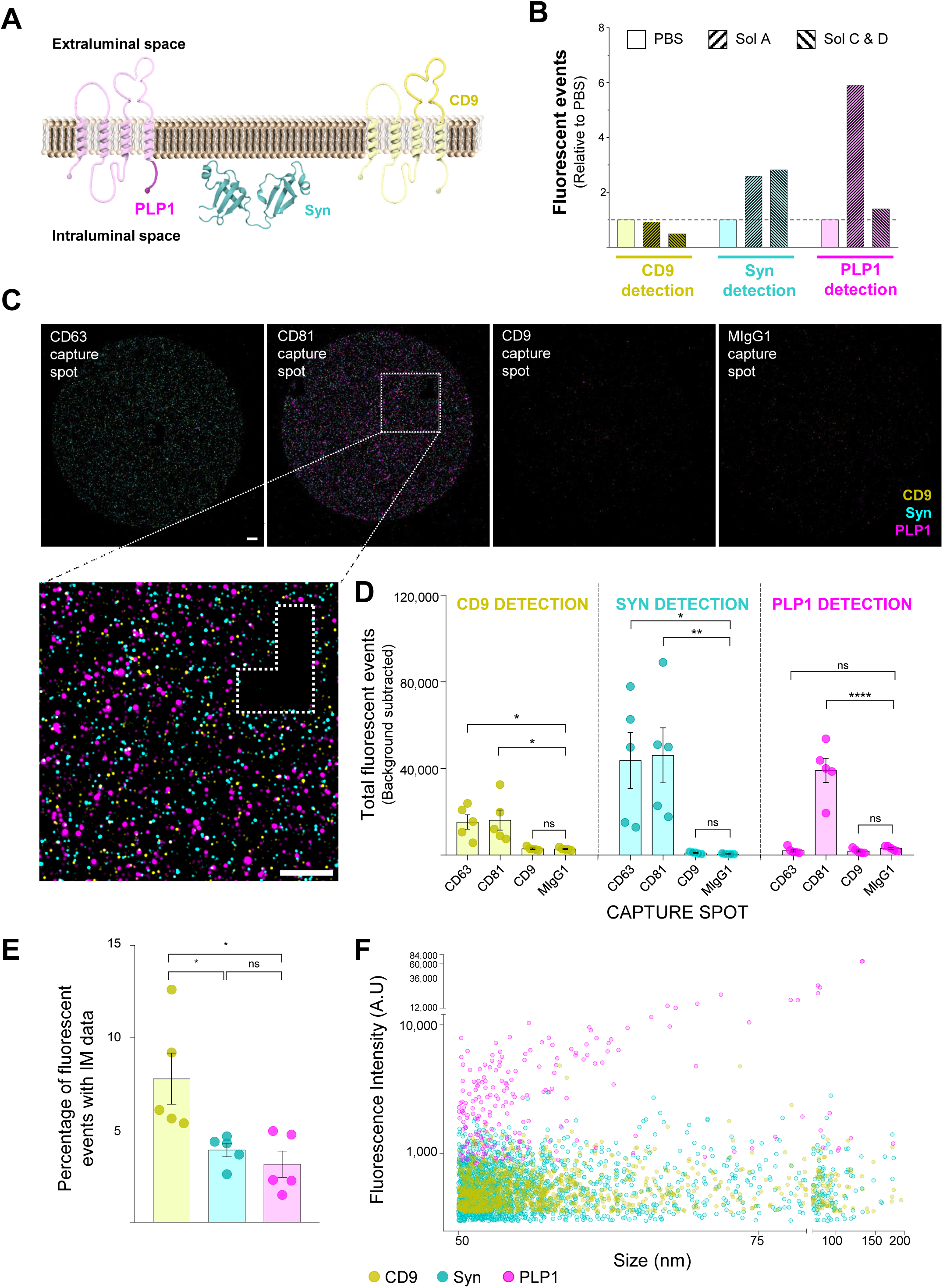
PLP1+ fluorescent events satisfy SP-IRIS/IF validation criteria but exhibit atypical fluorescence intensity. (**A**) Schematic of an EV lipid bilayer illustrating the relative orientations of target proteins PLP1 (magenta), Syn (cyan) and CD9 (yellow). Detection antibodies recognize an intraluminal epitope of PLP1, Syn and an extraluminal epitope of CD9. (**B**) Quantification of fluorescent events for CD9, Syn, and PLP1 detection antibodies across different buffer conditions relative to PBS in hNR-CM. Chips incubated with PBS (□), solution A (⎚) or solutions C and D (⎚). (**C**) Representative micrographs showing single-EV events using detection antibodies against CD9 (yellow), Syn (cyan) and PLP1 (magenta) in capture spots. Scale bar: 10 μm. Dashed lines in capture spots delineate inset. (**D**) Total background-subtracted fluorescent events for CD9, Syn and PLP1 across capture spots and the negative isotype control. Data represents biological replicates displayed with mean ± SEM from biological replicates (n=5). One-way ANOVA followed by Tukey post hoc test or Kruskal-Wallis followed by Dunn’s post hoc test for non-normally distributed data (* *p* < 0.05, ** *p* < 0.01, **** *p* < 0.0001, ns: not significant). (**E**) Percentage of fluorescent events (CD9, Syn, and PLP1) with matched interferometric (IM) data. Data represents biological replicates displayed with mean ± SEM from biological replicates (n=5). One-way ANOVA followed by Tukey post hoc test (* *p* < 0.05, ns: not significant). (**F**) Scatter plot of individual CD9+ (yellow), Syn+ (cyan) and PLP1+ (magenta) events showing fluorescence intensity (arbitrary units, A.U.) against particle size (nm).

### PLP1+ fluorescent events satisfy SP-IRIS/IF validation criteria but exhibit atypical fluorescence intensity

To validate the fluorescent signal from detection antibodies, the SP-IRIS/IF platform includes an internal quality control where chips lack capture antibodies. Detection antibodies against PLP1, Syn and CD9 showed no signal on antibody unprinted sections (white inverted “L” - Figure 2C). Moreover, fluorescent event counts for each protein (PLP1, Syn and CD9) were considered positive only when significantly higher than the signal on the mouse isotype control (MIgG1) spots. This analysis showed that CD9 and Syn were enriched in CD63 and CD81 capture spots, whereas PLP1 was exclusively present in CD81 spots (Figure 2D). Furthermore, the average percentage of fluorescent events with associated interferometry (IM) data was between 3.1% and 7.8% across all three markers. CD9+ fluorescent events with IM data were significantly higher than PLP1+ and Syn+, with no significant differences between PLP1+ and Syn+ (Figure 2E). In addition, size distribution analysis revealed values within the expected range for small EVs (50-200 nm, Figure 2F). These results suggest that PLP1 detection antibody showed similar IM and size distribution parameters to those previously validated (CD9 and Syn) by the manufacturer for SP-IRIS/IF analysis. Despite satisfying standard EV parameters, PLP1 fluorescence intensity distribution revealed a discrepancy with CD9+ and Syn+ fluorescent events. While the latter displayed values mostly below 2,000 arbitrary units (A.U.), PLP1 exhibited atypically higher intensities ranging from 1,000 A.U. up to 60,000 A.U. (Figure 2F). Considering that CD9 and Syn are proteins highly associated with EV that probably exhibit higher density than PLP1, this fluorescent signal pointed toward potential artifacts rather than authentic EV cargo.

### SP-IRIS/IF validation controls reveal the non-vesicular origin of the PLP1 signal

To discriminate whether PLP1 signal w EV-associated or non-vesicular, we analyzed CD81 capture spots (where PLP1+ events are detected) across serial dilutions. While total Syn+ and CD9+ events decreased with CM dilution, PLP1 signal was not affected (Figure 3A-B). Moreover, blank (no sample) and detergent control (1% Triton X-100) showed no significant reduction in total PLP1+ counts (Figure 3C). Finally, no significant differences were observed among the blank, CM, and 1% Triton X-100 conditions regarding either the percentage of PLP1+ events with IM data or the overall fluorescence intensity distribution. In contrast, almost all Syn+ and CD9+ events were depleted in both control conditions (Figure 3D-E, see also Figure 2F). Collectively, our results indicate that although PLP1+ events satisfy most internal validation criteria, they do not originate from authentic EV cargo.

**Figure 3.**
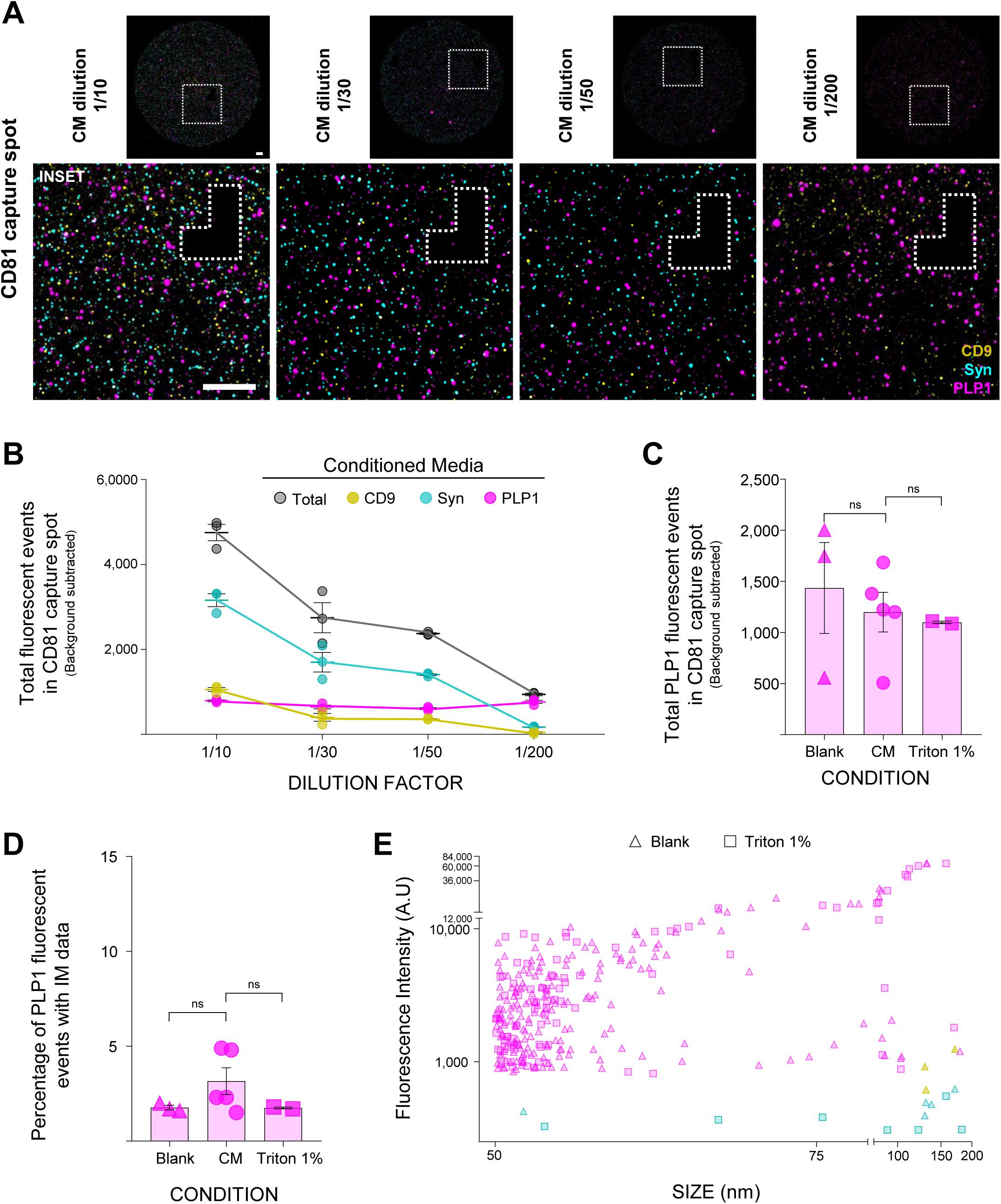
SP-IRIS/IF validation controls reveal the non-vesicular origin of the PLP1 signal. (**A**) Representative micrographs showing single-EV events using detection antibodies against CD9 (yellow), Syn (cyan) and PLP1 (magenta) in CD81 capture spots across serial dilutions (1/10, 1/30, 1/50, and 1/200) of hNR-CM. Dashed lines in capture spots delineate antibody unprinted sections. Scale bar: 10 μm. (**B**) Total background-subtracted fluorescent events on CD81 capture spots plotted against dilution factor for total events (gray), CD9 (yellow), Syn (cyan), and PLP1 (magenta). Data represent technical replicates displayed with mean ± SEM. (**C**) Total background-subtracted PLP1+ events on CD81 capture spots in control without sample (Blank), hNR-CM (CM) and hNR-CM with 1% Triton X-100 (Triton 1%). Data represents biological replicates displayed with mean ± SEM from biological replicates (n=2-5). Kruskal-Wallis followed by Dunn’s post hoc test (ns: not significant). (**D**) Percentage of detected PLP1+ events with matched IM data in control without sample (Blank), hNR-CM (CM) and hNR-CM with 1% Triton X-100 (Triton 1%). Data represents biological replicates displayed with mean ± SEM from biological replicates (n=2-5). Kruskal-Wallis followed by Dunn’s post hoc test (ns: not significant). (**E**) Scatter plot of individual CD9+ (yellow), Syn+ (cyan) and PLP1+ (magenta) events showing fluorescence intensity (arbitrary units, A.U.) against particle size (nm) under Blank (triangles) and 1% Triton X-100 (squares) conditions.

### PLP1 fluorescent artifact is independent of sample matrix and sensor chip layouts

To determine whether the observed artifact was dependent on the biological matrix or the chip layout, we analyzed human serum samples using ExoView Plasma chips. These include an additional capture spot for platelet-derived EVs (CD41a+) alongside tetraspanins. Consistent with our findings in CM, EVs associated signals (CD81+ and CD9+ fluorescent events) were efficiently depleted following 1% Triton X-100 lysis and remained undetectable in blank controls across all capture spots in two serum samples. Although PLP1+ event counts on CD81 capture spots showed a variable response across samples, unlike CD81+ and CD9+ events, they did not completely disappear upon detergent treatment and persisted at high levels in the blank control (Figure 4A-B). To test if PLP signal was specific to detection antibodies or generated by capture layouts, we functionalized ExoView Flex chips with an anti-PLP1 antibody (Figure 4C). Using this customized layout, we confirmed that PLP1+ fluorescent events on CD81 capture spots behaved similarly to those on the standard chips previously used. In the PLP1 FLEX capture spots, blank controls did not show any fluorescent events in all three serum samples used. However, when sera were incubated on the chips, PLP1+ events appeared but completely lacked EV markers (CD81 and CD9). Furthermore, PLP signal was not disrupted by 1% Triton X-100, resulting in highly variable particle counts across samples (Figures 4D–E). These results indicate that the non-vesicular PLP1+ events are independent of the biological matrix (conditioned media and serum) or the sensor chip configuration (tetraspanin, plasma or Flex chips). In summary, our data underscores the importance of rigorous controls for single-EV platforms such as SP-IRIS/IF, highlighting detergent-based membrane lysis as the gold standard for true EV validation.

**Figure 4.**
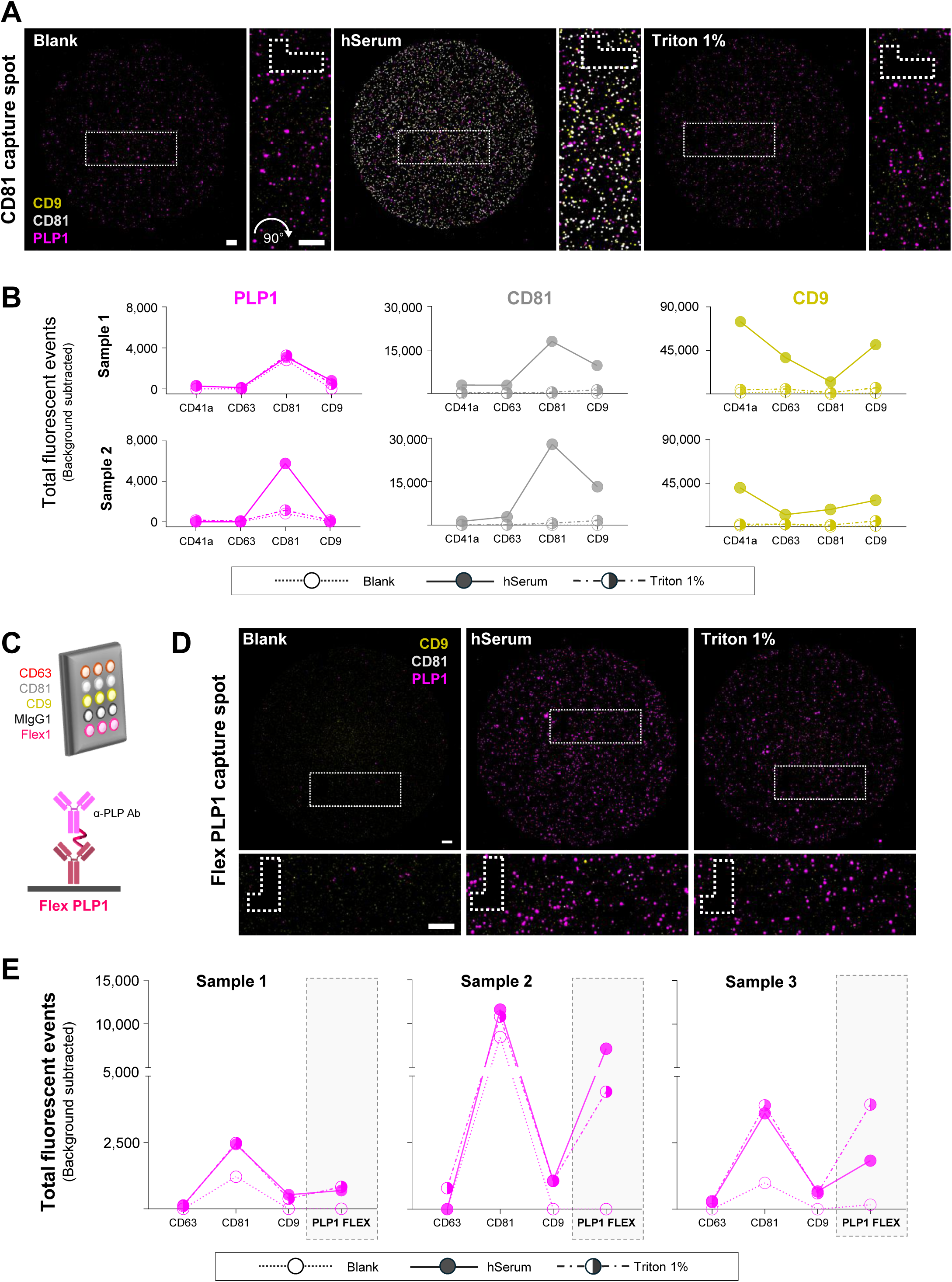
PLP1+ fluorescent artifact is independent of sample matrix and sensor chip layouts. (**A**) Representative micrographs showing single-EV events using detection antibodies against PLP1 (magenta), CD81 (gray) and CD9 (yellow) in CD81 capture spots in control without sample (Blank), human serum sample (hSerum), and human serum treated with 1% Triton X-100 (Triton 1%). The inset image has been rotated 90° clockwise relative to the parent field to optimize panel layout. Scale bar: 10 μm. (**B**) Total background-subtracted fluorescent events quantified on CD41a, CD63, CD81 and CD9 capture spots using detection antibodies against PLP1 (magenta), CD81 (gray) and CD9 (yellow) for Sample 1 and Sample 2. Blank (dotted line), hSerum (solid line), and Triton 1% (dashed-dotted line). (**C**) Schematic depicting Flex sensor chip layout using customized spots functionalized with an anti-PLP1 antibody (Flex PLP1). (**D**) Representative micrographs showing single-EV events using detection antibodies against PLP1 (magenta), CD81 (gray) and CD9 (yellow) in Flex PLP1 capture spots in control without sample (Blank), human serum sample (hSerum), and human serum treated with 1% Triton X-100 (Triton 1%). Scale bar: 10 μm. (**E**) Total background-subtracted PLP1+ events quantified on CD63, CD81, CD9 and Flex PLP1 capture spots for Sample 1, Sample 2 and Sample 3. Blank, (dotted line), hSerum (solid line), and Triton 1% (dashed-dotted line).

## Discussion

SP-IRIS coupled with immunofluorescence (SP-IRIS/IF) relies on the selective capture of EVs, where single-EV size and count are quantified via interferometry, while the molecular phenotype of captured EVs is profiled by detection antibodies (Daaboul et al., 2011, 2016). In this work, by evaluating a detection antibody across different biological matrices and sensors, we uncovered non-vesicular signals capable of generating both interferometric and fluorescent signals that closely mimic EV parameters. Our findings emphasize that detergent lysis controls are critical for recognizing authentic single-EV signal in SP-IRIS/IF. Particularly, these controls are critical when introducing antibodies or probes not previously validated at single-particle resolution. Altogether, our data highlights the importance of including lysis controls, especially in single-EV platforms currently considered as promising tools for biomarker discovery and translational research.

The 2023 MISEV guidelines recommend including vesicle-disrupting surfactants (detergents) controls to confirm the association of fluorescent events to intact lipid membranes for single-EV platforms such as SP-IRIS/IF (Welsh et al., 2024). Although previous works have shown their importance in other single-particle techniques (Görgens et al., 2019; György et al., 2011; Welsh et al., 2020), the routine implementation of these controls remains surprisingly sparse in the field. Here we found that non-vesicular signals can closely resemble EV parameters, indicating that SP-IRIS/IF analyses are equally susceptible and can lead to data misinterpretation. Although these results were obtained by using antibodies targeting a specific EV cargo, detergent-mediated lysis should be considered mandatory for single-EV analysis.

The myelin proteolipid protein (PLP1) has been extensively validated as an EV cargo in different cellular and animal models using a diverse set of techniques (Frühbeis et al., 2020; Herrera Lopez et al., 2025; Krämer-Albers et al., 2007). Therefore, we aimed to use SP-IRIS/IF to study its distribution at the single-EV level. Our findings highlight a critical technical limitation regarding antibody-based detection on this platform. Despite following quality controls, we detected a non-vesicular artifact that closely mimics true EV cargo and whose molecular origin remains unclear. The detection antibody used against PLP1 targets a short linear epitope (residues 272-277), which increases the probability of cross-reactivity with capture antibodies. Nevertheless, the association of fluorescent events with IM data (size) and the artifact’s persistence on FLEX chips recognizing a longer epitope of PLP1 (residues 177-232), suggest a more complex interpretation that needs further studies.

While SP-IRIS/IF optimizations could explore different concentrations or alternative surfactants, 1% Triton X-100 is a frequently implemented strategy in single-EV analysis (Bravo-Miana et al., 2024; Görgens et al., 2019; Welsh et al., 2024). Importantly, this concentration effectively decreased the signal of EV-associated proteins (Syn, CD9 and CD81), suggesting successful lipid membrane disruption while confirming the non-vesicular nature of the PLP1 signal. Although assessing whether this signal is reproduced across other single-EV platforms falls beyond the scope of our study, it represents a valuable consideration for the broader EV field. Despite these limitations, this report establishes a practical case that highlights the importance of applying robust methodological practices to study EVs and differentiating them from other non-vesicular particles (György et al., 2011; Jeppesen et al., 2019; Sódar et al., 2016) or antibody-mediated artifacts.

The EV field has expanded rapidly, transitioning from descriptive cell biology into clinical trials and high-throughput diagnostic platforms. However, this progress raises a critical question: what operationally defines an EV? At single-particle resolution, vesicular status is usually inferred via indirect measurements that combine optical methods with affinity-based fluorescent probes. As shown here, non-vesicular signals can fully satisfy these operational size and intensity gates, creating false positives indistinguishable from authentic vesicles. Because the hallmark of an extracellular vesicle is an intact lipid bilayer (Welsh et al., 2024), analytical validity demands a biochemical demonstration. Therefore, detergent lysis controls are essential to prevent data misinterpretation and ensure that single-EV platforms rest on verifiable physical foundations.

## Acknowledgements

The authors thank Dr. Alfredo Cáceres and Dr. Laura Morelli for providing hiPSCs culture lines. Also, Dr. Mónica Remedi and Jone Karmele Arizaga Echevarría for their technical assistance and training with cell culture and single-EV analysis, respectively.

## Author Contributions

**G.G.G.:** Conceptualization, Formal Analysis, Funding Acquisition, Investigation, Methodology, Visualization, Writing – Original Draft**; D.O.:** Conceptualization, Funding Acquisition, Supervision, Writing – Review & Editing; **H.C.-V.:** Investigation, Project Administration, Resources, Writing – Review & Editing**; D.C.-C.:** Resources, Writing – Review & Editing; **R.C.B.-M.:** Conceptualization, Funding Acquisition, Methodology, Supervision, Writing – Review & Editing; **A.L.M.:** Conceptualization, Funding Acquisition, Supervision, Visualization, Writing – Original Draft.

## Data Availability

The data that supports the findings of this study are available from the corresponding author upon reasonable request.

